# Ecological and life-history traits and their relationship with West Nile virus and Saint Louis Encephalitis virus exposure risk

**DOI:** 10.1101/2023.01.30.526345

**Authors:** O. Giayetto, A.P. Mansilla, F.N. Nazar, A. Diaz

## Abstract

Host life-history traits can influence host-vector encounter rates, and so differentially determine the exposure risk of bird species. This modulation of host-virus encounters’ dynamics is especially important when facing “generalist” arboviruses like West Nile virus (WNV) and Saint Louis Encephalitis virus (SLEV). Using prevalence data collected by our laboratory since 2004, we tested several hypothesis that included birds’ ecological and life-history traits to determine which traits were better predictors of birds’ exposure risk to these arboviruses. By means of information-theoretic procedures and generalized mixed linear models, we observed that body mass was an important trait when predicting birds’ exposure risk to WNV and SLEV and migratory status significantly influenced birds’ exposure risk only to WNV. Our study highlights important traits to consider when studying the transmission system of these arboviruses, being useful to focus resources when characterizing viral transmission networks and discuss the repercussions of these traits over birds’ immune function throughout the pace of life syndrome and trade-offs theory.

## Introduction

West Nile Virus (WNV) and Saint Louis Encephalitis Virus (SLEV) are two emerging arboviruses belonging to the family Flaviviridae. Both are causative agent of encephalomyelitis and are considered of global health concern (Guth et al. 2020), affecting not only humans but also wildlife (LaDeau et al. 2007; Ong et al. 2021). Both, WNV and SLEV are maintained in a diverse ‘multi-vector multi-host’ transmission network, with intrinsic spatial and seasonal fluctuation (Diaz et al. 2013b). West Nile and SLE viruses share an ecological resemblance in their maintenance network being predominantly maintained by members of the Culex genera and a wide range of avian hosts’ species (mainly Passeriformes and Columbiformes) (Reisen 2003, 2013; Diaz et al. 2013a; Ciota 2017; Rochlin et al. 2019; Giayetto et al. 2021). However, there are differences in their hosts’ species range and in the consequences that these viruses might have on the health status of their hosts. For example, WNV tends to have an increased mortality in a higher range of bird species (Lanciotti et al. 1999; Lord & Day 2001; Maharaj et al. 2018).

To be maintained in nature, both WNV and SLEV need a competent vector to feed on a competent host. Consequently, mosquitoes feeding patterns take an important role in viral dynamics (Kilpatrick et al. 2006a), affecting host-vector encounter rates and favouring certain host species to be chosen by competent vectors. Despite certain evolutionary and environmental conditions that affect mosquitos feeding patterns-i.e., host availability, hosts defensive behaviours and blood nutritive value- there are interspecific differences in host life-history traits that can influence host-vector encounter rates, and so differentially determine the exposure risk of bird species and modulate the host-viral encounters dynamics (Lyimo & Ferguson 2009; Kernbach et al. 2021).

Some of the life history traits that can have repercussions in modelling exposure risk in host species are nest characteristics, body mass, clutch size, migratory behaviour, and diet type, among others. Particularly, it has been described nest height and type were able to predict the infection probability to heamosporadian parasites transmitted by Culex species in central and southern African birds (Lutz et al. 2015; Ganser et al. 2020). Body mass and clutch size affect vector-hosts interactions since larger and bigger bodies as well as bigger clutches releases more olfactory cues to vectors, as observed by Figuerola et al. where a greater seroprevalence of WNV was observed in larger hosts (Figuerola et al. 2008; Takken & Verhulst 2013). Migratory behaviour and seasonality also influence vector-host encounters by affecting mosquito host availability (Rappole et al. 2000; Peterson et al. 2004). Finally, differences in hosts’ diet type have previously been associated with the infection risk to other arboviruses, potentially due to the relationship between the type of diet and the foraging time during which an individual may be exposed to a mosquito bite (González et al. 2014; Walsh 2019; Skinner et al. 2021).

In the previously described frame, we hypothesize ecological and life history traits can modulate vector-host encounters and consequently predict birds’ exposure risk to vector-borne pathogens, we expect differences among traits when trying to predict exposure risk to WNV and SLEV in birds, with some of such traits being better predictors than others. In evaluating whether the selected ecological and life history traits are able to predict birds’ exposure risk to WNV and SLEV, we were able to establish that certain hypothesis (and consequentially, certain traits) are better when predicting birds’ exposure risk to WNV and SLEV.

## Methods

### Prevalence data

We used the serological state against WNV and SLEV (presence/absence of neutralizing antibodies) data from free-ranging birds, obtained between 2004 and 2019 in 5 different Argentinean provinces (Córdoba, Chaco, La Pampa, San Luis, and Tucumán) (Figure 1). For Córdoba province, we used data obtained in 2004, 2005, 2006, 2012, 2013 and 2014 from Cordoba city, and La Para (Diaz et al. 2008, 2016). For Chaco and Tucumán provinces, data were obtained in 2005 and 2006 respectively (Diaz et al. 2008). The province of San Luis was sampled in 2011. Finally, data from La Pampa was obtained between 2017 and 2019 (Mansilla et al. 2022). Within each province, several sample sites were used (Figure 1). The presence of neutralizing antibodies in avian sera was used as indicative of a previous exposure to WNV and/or SLEV and the posterior survival of the individual. Birds’ sampling procedure was performed as described elsewhere (Diaz et al. 2016). Briefly, birds were captured using mist nets. After capture, birds were taxonomically determined and then bleed by jugular veinpuncture. Blood samples were diluted in Minimum essential Medium and then centrifuged for serum obtention. Serum was stored at −20°C until analysis. Finally, samples were screened for the presence of WNV- and SLEV-reactive antibodies by plaque-reduction neutralization test (Tauro et al. 2012).

**Figure 1.**
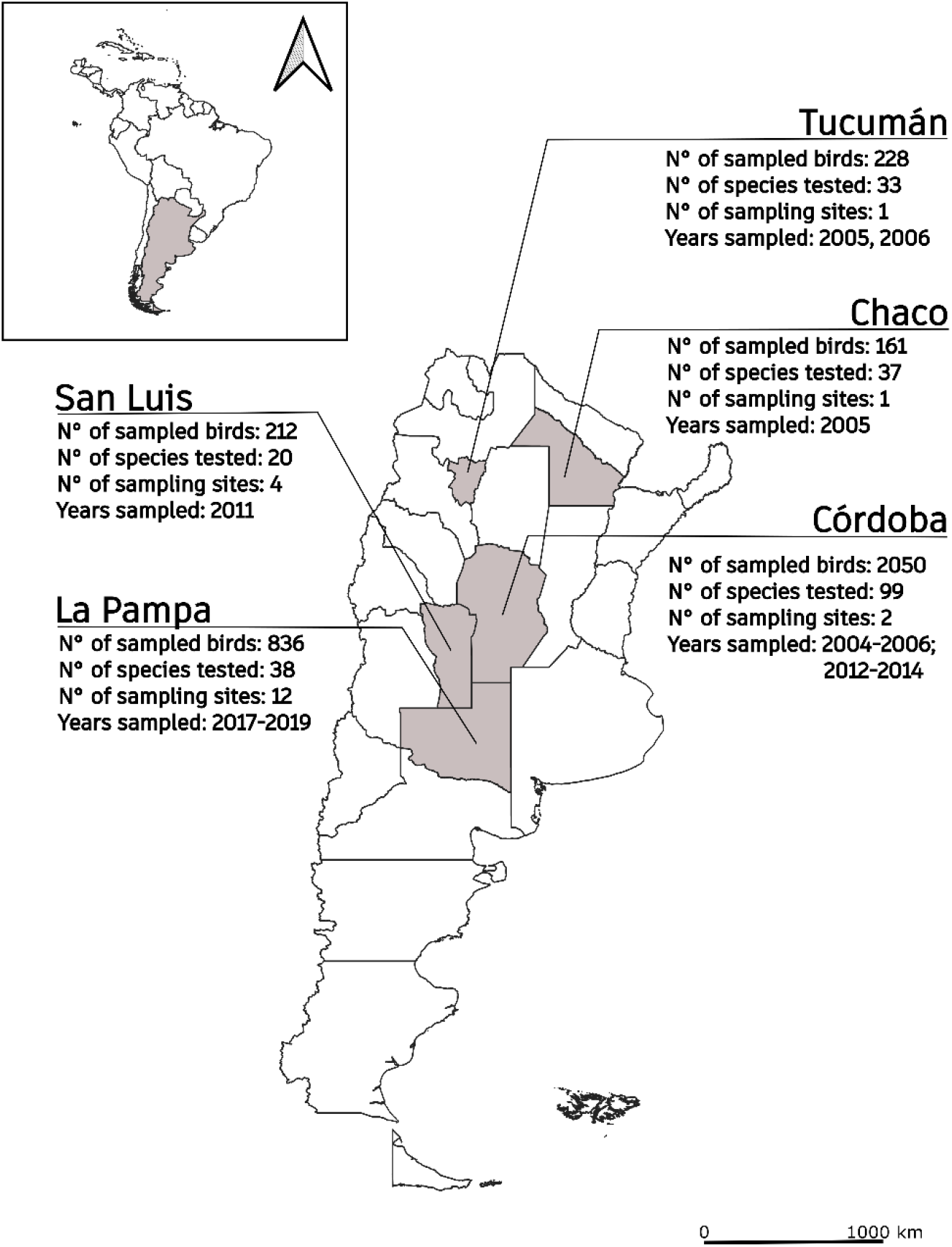
Map of sampled Argentinean provinces. Within each province, the number of sampled birds, species tested, sample sites and years sampled are included.

### Ecological and life-history traits data

We combined a dataset of ecological and life-history traits that were potentially relevant to explain WNV and SLEV birds’ exposure risk (Rappole et al. 2000; Peterson et al. 2004; Figuerola et al. 2008; Takken & Verhulst 2013; González et al. 2014; Lutz et al. 2015; Walsh 2019; Ganser et al. 2020; Skinner et al. 2021). We selected the following six traits: body mass, clutch size, nest height, nest type, diet type and migratory status. Both, body mass and clutch size were included as each species’ media value. Nest height was classified in three categories, as ground, understory (nests predominantly found above ground and below 3m) or canopy/sub-canopy (nests predominantly found above 3m). Nest type was classified in three categories, as open cup, closed cup or cavity. Diet type was categorized into 7 groups according to the primary diet of each species: Folivore-Frugivore, Frugivore-Insectivore, Granivore, Granivore-Insectivore, Insectivore, Insectivore-Carnivore and Omnivore. Finally, migratory status was categorized as resident or migrant. For body mass and clutch size, data was extracted from Cooke et al., 2019 dataset (Data available from figshare repository: https://doi.org/10.6084/m9.figshare.5616424.v1) (Cooke et al. 2019). Information referring to nest characteristics, diet and migratory status was extracted mainly from The Cornell Lab of Ornithology website (https://birdsoftheworld.org). We supplemented missing species traits with additional data to have all the mentioned traits for each species included in this study (Mason 1986; Wiley 1988; Aráoz et al. 2016).

### Statistical analysis

To test the effects of ecological and life-history traits on birds’ exposure risk, data including WNV and SLEV birds’ serological status was used as response variable (Kernbach et al. 2021). We used generalised linear mixed models (GLMM) with Binomial error distribution and logit link function. Nest type, nest height, migratory status, migration type and diet type were included as categorical fixed effects, while clutch size and body mass were standardized (mean=0; SD=1) and included as continuous fixed effects. As both viruses shown stational and geographical variations in their circulation (Diaz et al. 2013b), sampled year and sample sites nested within province were included as a random effect. Birds’ family was also included as a random effect to account for the statistical lack of independence within host phylogenetic relationships.

Twenty candidate models for each virus were evaluated following information-theoretic procedures (Burnham & Anderson 2002). We considered models that tested each of our candidate hypothesis over ecological and life history traits that could account for birds’ exposure risk to SLEV and WNV (Table 1). Within the candidate models, we included those with the individual effects of each of the selected traits as well as models containing the additive effects of certain mentioned traits.

**Table 1.**
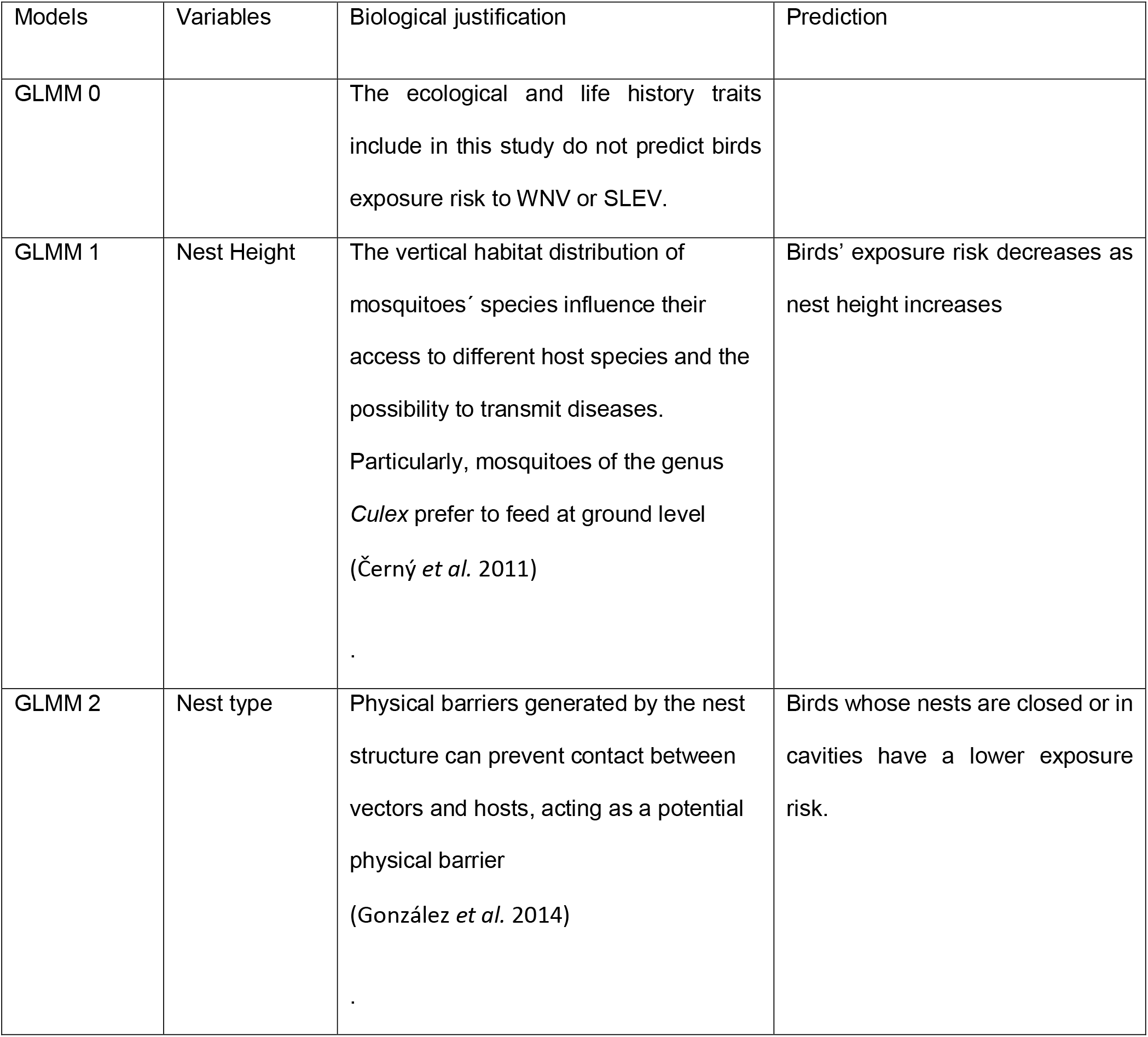
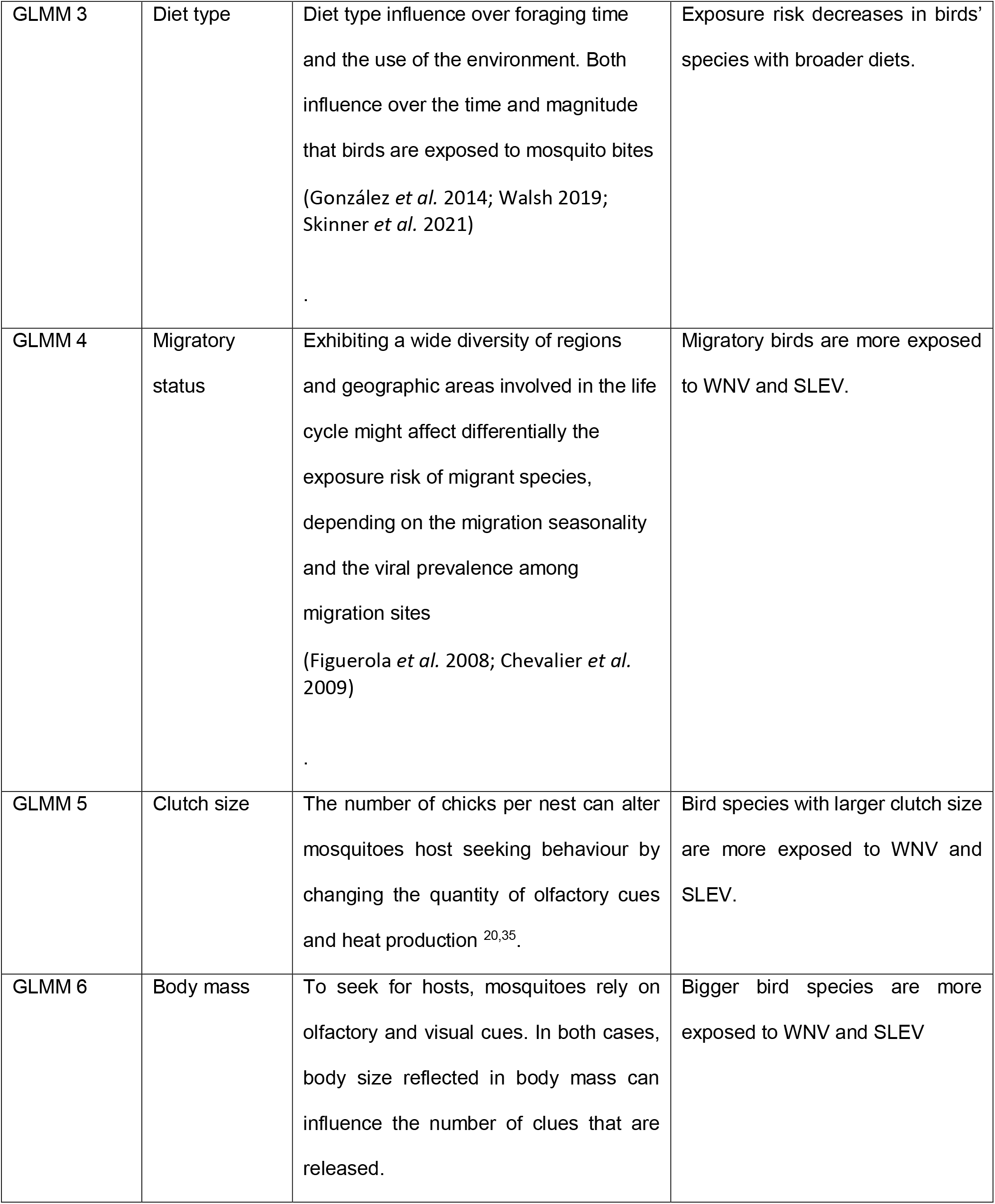
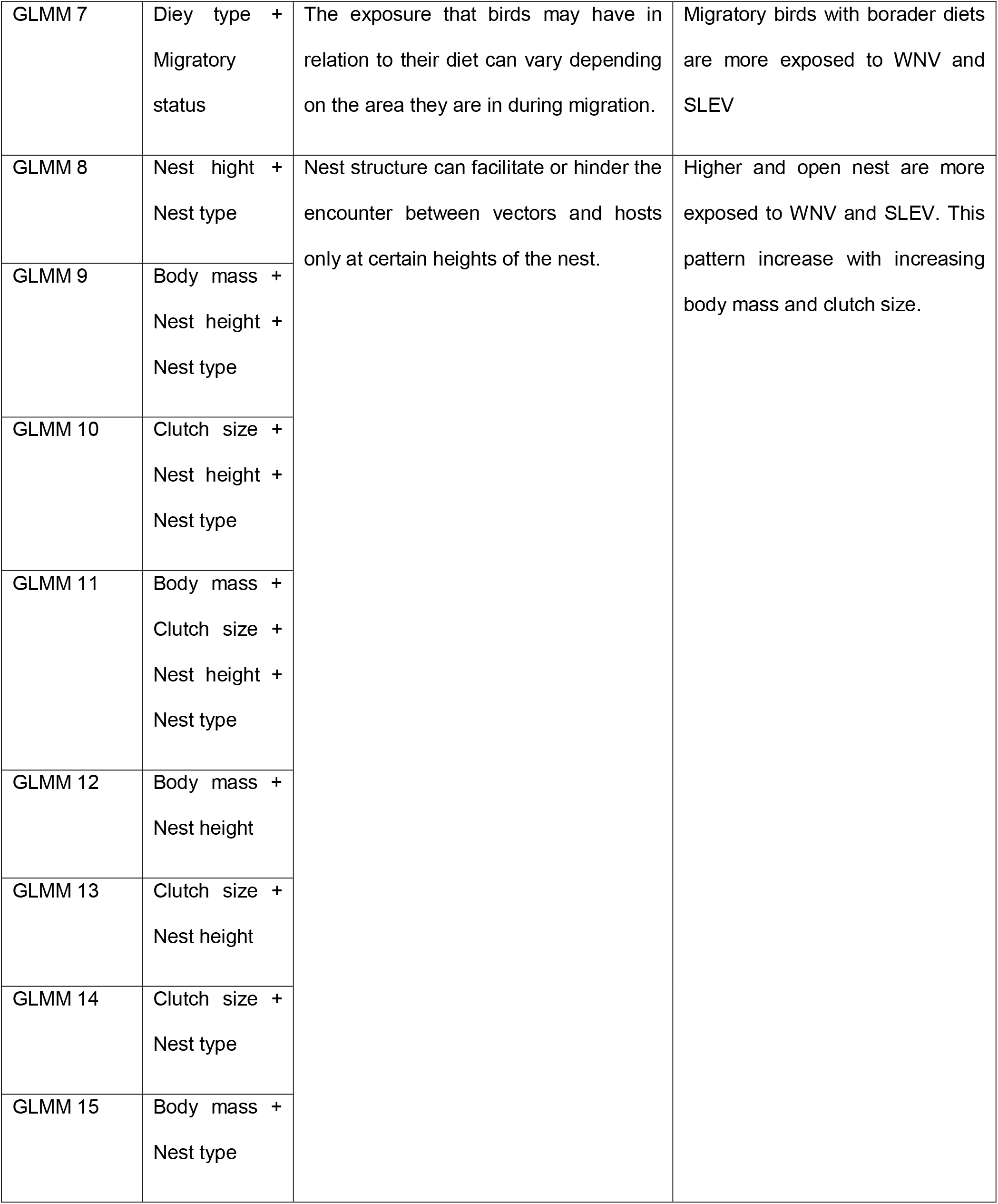
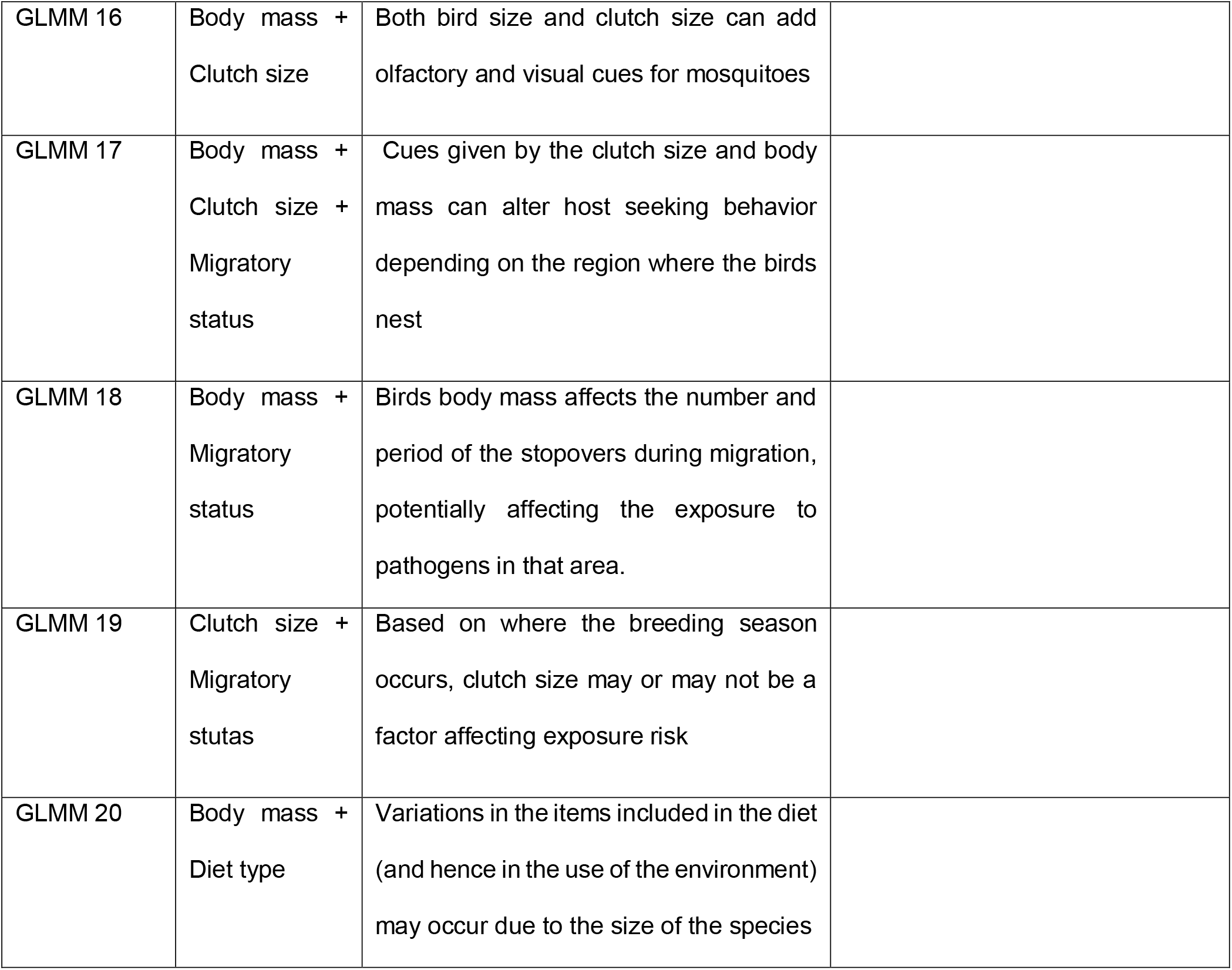
Structure of Generalized Linear Mixed Models analysing the association between birds’ exposure risk to West Nile Virus (WNV) or Saint Louis Encephalitis Virus (SLEV) and ecological and life history traits. For all models, sampled year, sample sites nested within province and birds’ family were used as random effects. For each model, the biological justification and predictions are explicated.

| Models | Variables | Biological justification | Prediction |
| --- | --- | --- | --- |
| GLMM 0 |  | The ecological and life history traits include in this study do not predict birds exposure risk to WNV or SLEV. |  |
| GLMM 1 | Nest Height | The vertical habitat distribution of mosquitoes' species influence their access to different host species and the possibility to transmit diseases. Particularly, mosquitoes of the genus *Culex* prefer to feed at ground level (Černý *et al.* 2011) | Birds' exposure risk decreases as nest height increases |
| GLMM 2 | Nest type | Physical barriers generated by the nest structure can prevent contact between vectors and hosts, acting as a potential physical barrier (González *et al.* 2014) | Birds whose nests are closed or in cavities have a lower exposure risk. |
| GLMM 3 | Diet type | Diet type influence over foraging time and the use of the environment. Both influence over the time and magnitude that birds are exposed to mosquito bites (González *et al.* 2014; Walsh 2019; Skinner *et al.* 2021) | Exposure risk decreases in birds' species with broader diets. |
| GLMM 4 | Migratory status | Exhibiting a wide diversity of regions and geographic areas involved in the life cycle might affect differentially the exposure risk of migrant species, depending on the migration seasonality and the viral prevalence among migration sites (Figuerola *et al.* 2008; Chevalier *et al.* 2009) | Migratory birds are more exposed to WNV and SLEV. |
| GLMM 5 | Clutch size | The number of chicks per nest can alter mosquitoes host seeking behaviour by changing the quantity of olfactory cues and heat production <sup>20,35</sup>. | Bird species with larger clutch size are more exposed to WNV and SLEV. |
| GLMM 6 | Body mass | To seek for hosts, mosquitoes rely on olfactory and visual cues. In both cases, body size reflected in body mass can influence the number of clues that are released. | Bigger bird species are more exposed to WNV and SLEV |
| GLMM 7 | Diet type +<br>Migratory status | The exposure that birds may have in relation to their diet can vary depending on the area they are in during migration. | Migratory birds with broader diets are more exposed to WNV and SLEV |
| GLMM 8 | Nest height +<br>Nest type | Nest structure can facilitate or hinder the encounter between vectors and hosts only at certain heights of the nest. | Higher and open nest are more exposed to WNV and SLEV. This pattern increase with increasing body mass and clutch size. |
| GLMM 9 | Body mass +<br>Nest height +<br>Nest type |  |  |
| GLMM 10 | Clutch size +<br>Nest height +<br>Nest type |  |  |
| GLMM 11 | Body mass +<br>Clutch size +<br>Nest height +<br>Nest type |  |  |
| GLMM 12 | Body mass +<br>Nest height |  |  |
| GLMM 13 | Clutch size +<br>Nest height |  |  |
| GLMM 14 | Clutch size +<br>Nest type |  |  |
| GLMM 15 | Body mass +<br>Nest type |  |  |
| GLMM 16 | Body mass +<br>Clutch size | Both bird size and clutch size can add olfactory and visual cues for mosquitoes |  |
| GLMM 17 | Body mass +<br>Clutch size +<br>Migratory<br>status | Cues given by the clutch size and body mass can alter host seeking behavior depending on the region where the birds nest |  |
| GLMM 18 | Body mass +<br>Migratory<br>status | Birds body mass affects the number and period of the stopovers during migration, potentially affecting the exposure to pathogens in that area. |  |
| GLMM 19 | Clutch size +<br>Migratory<br>status | Based on where the breeding season occurs, clutch size may or may not be a factor affecting exposure risk |  |
| GLMM 20 | Body mass +<br>Diet type | Variations in the items included in the diet (and hence in the use of the environment) may occur due to the size of the species |  |

Models were tested for Akaike’s information criterion corrected for small sample size (AICc) (Burnham & Anderson 2002, 2004) for the analysis of variation in WNV and SLEV seroprevalence. Model comparisons were made with ΔAICc, calculated as the difference in the AICc values between each model and the best-fitted model in the set. Akaike weight (ω AICc) was also calculated for each model to evaluate the weight of evidence in favour of each candidate model. Parameter estimates were calculated using model-averaged parameter estimates based on the ω AICc from a subset of candidate models. The subset was stablished considering models with ΔAICc < 7 (Burnham & Anderson 2004). To supplement parameter-likelihood evidence of important effects, we calculated 95% confidence interval limits (CI) of parameter estimates, considering that a parameter is significantly associated with the WNV or SLEV exposure risk when its CI did not include zero. Statistical analyses were performed in the ‘lme4’ package and ‘MuMIn’ (Bates et al. 2015; Barton 2020), in R v 4.0.3 (R Core Team, 2020). For data visualisation of birds seroprevalence we used non-metric multidimensional scaling (k=2) using Gower similarities distance between species ordered by their ecological and life history traits with the ‘metaMDS’ function from the vegan package (Oksanen et al. 2020) (Figure 2).

**Figure 2.**
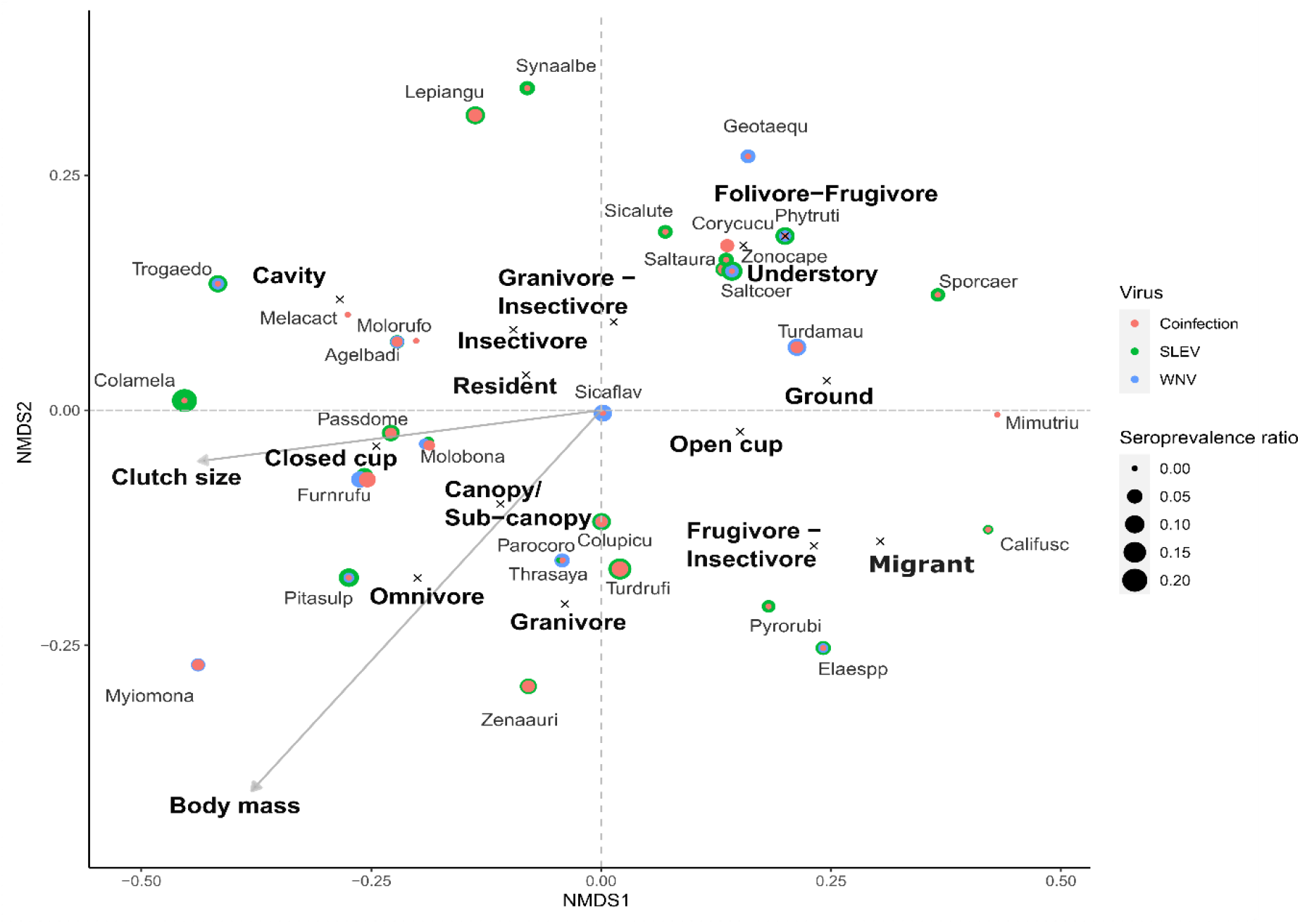
Visualization of birds’ species according their seroprevalence and their ecological and life history traits. Non-metric multidimensional scaling (NMDS) plot of life history and ecological traits (via Gower dissimilarities distance) between sampled species with more than 20 individuals tested for West Nile virus (WNV) and Saint Louis Encephalitis virus (SLEV) neutralizing antibodies (n = 33). Concentric circles show if the evaluated species included individuals that were positive for WNV, SLEV antibodies or showed coinfections between both viruses and the circle size indicates the magnitude of the seroprevalence.

## Results

A total of 3487 tested birds were included in our analysis. WNV seroprevalence accounted for 4% distributed in 42 species. On the other hand, 8% of the tested birds were positive to SLEV distributed in 60 species. Only 1.5% of the avian sera tested performed a heterotypic immunological reaction indicating birds had been exposed to both viruses (Figure 2). As a first indicative of virus-host contact, birds belonging to Bucconidae, Caprimulgidae, Polioptilidae families where only exposed to WNV, whereas birds belonging to Charadriidae, Emberizidae, Falconidae, Mimidae, Passerellidae, Picidae, Recurvirostridae and Scolopacidae families where only exposed to SLEV.

Among the 20 models tested for both WNV and SLEV, 10 models were selected (ΔAICc <7) for WNV and 7 for SLEV that could predict birds’ exposure risk to these viruses (Table 3 and 4). The model-averaged for WNV included migratory status, body mass, clutch size, nest height, diet type and nest type as explanatory variables. The 95% CI for the parameter estimates for migratory status (0.85 relative importance) and body mass (0.84 relative importance) did not include zero, indicating that these two explanatory variables significantly influenced birds’ exposure risk to WNV (Figure 3). West Nile virus exposure risk increased 1.22 times for each increase in the standard deviation (28.6 g) of body mass (0.196 ± 0.07, estimate and SD) and differed between migratory status, being resident birds 5 times more exposed than migrants (−2.89 ± 1.5; −4.47 ± 0.88, estimates and SD for residents and migrant birds, respectively) (Figure 3). For SLEV, the model-averaged included migratory status, body mass, clutch size, nest height, and nest type as explanatory variables. The 95% CI for the parameter estimates for body mass (one relative importance) did not include zero, indicating that this explanatory variable significantly influenced birds’ exposure risk to SLEV (Figure 3). SLEV exposure risk was positively associated with body mass, increasing 1.18 times for each increase in the standard deviation (28.6 g) of body mass (0.176 ± 0.06, estimate and SD).

**Figure 3.**
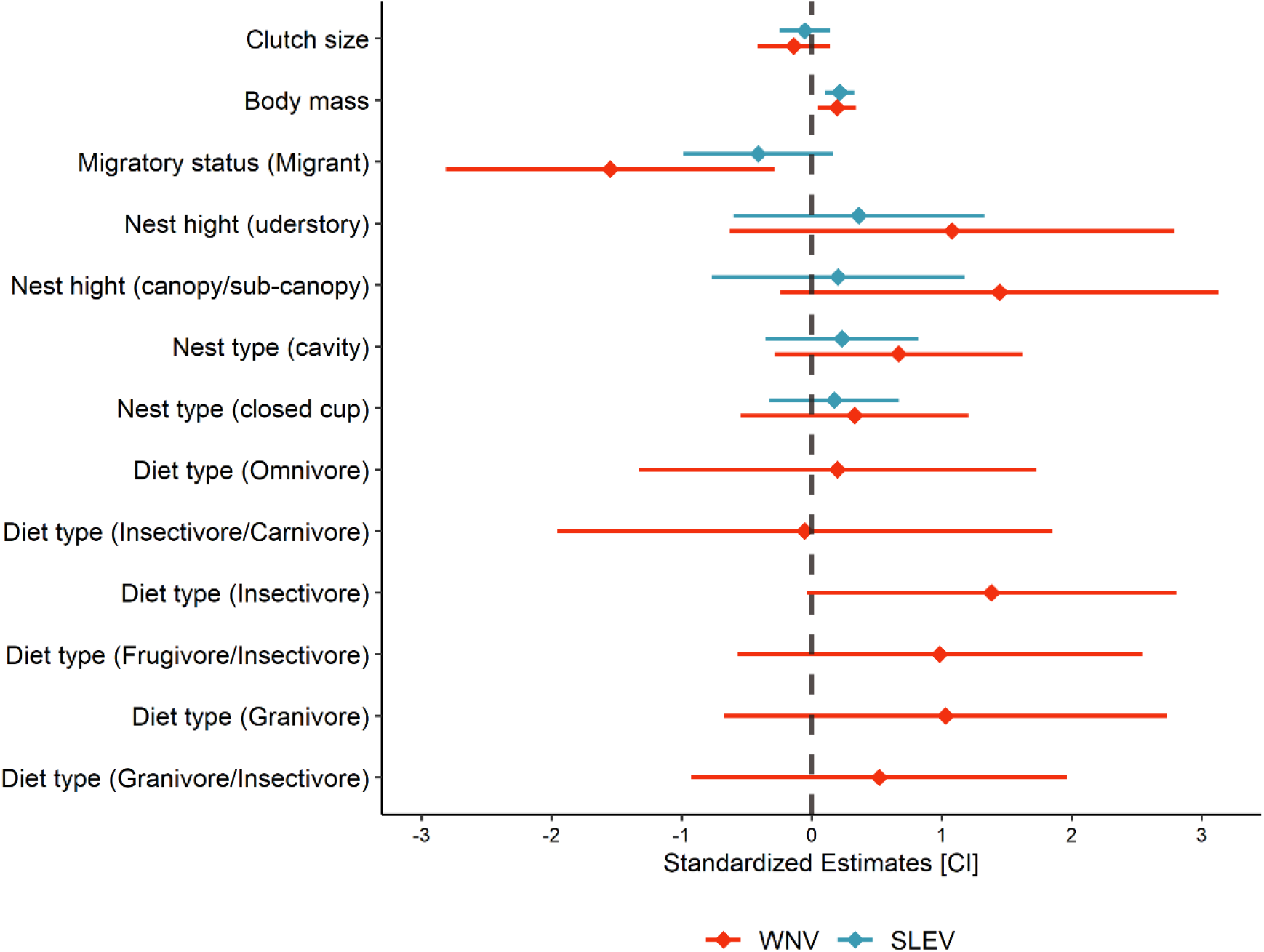
Effect sizes of the selected ecological and life-history traits variables on West Nile virus (red) and Saint Louis Encephalitis virus (blue) exposure risk. Standardized effect sizes (rhombus symbols) and 95% CI (lines) from model-averaged parameter estimates from the subset of candidate models. Body mass and migratory status had significative effects on WNV risk, whereas only body mass had significative effects on SLEV risk. In both cases, body mass is positively associated, and for WNV, residents were more exposed than migrants were.

**Table 2.** Summary of model-selection results for models explaining variation in WNV exposure risk. k is the number of estimated parameters. Models are listed in decreasing order of importance. Models selected for model-averaged with ΔAICc > 7 are in bold.

| Candidate Models | k | AICc | $\Delta AICc$ | $\omega AICc$ |
| --- | --- | --- | --- | --- |
| **GLMM 18** | **6** | **1071.5** | **0.00** | **0.420** |
| **GLMM 17** | **7** | **1072.6** | **1.11** | **0.242** |
| **GLMM 4** | **5** | **1074.6** | **3.12** | **0.088** |
| **GLMM 19** | **6** | **1075.6** | **4.14** | **0.053** |
| **GLMM 12** | **7** | **1075.7** | **4.25** | **0.050** |
| **GLMM 6** | **5** | **1076.1** | **4.62** | **0.042** |
| **GLMM 16** | **6** | **1077.6** | **6.10** | **0.020** |
| **GLMM 7** | **11** | **1077.9** | **6.46** | **0.017** |
| **GLMM 11** | **10** | **1078.3** | **6.78** | **0.014** |
| **GLMM 9** | **9** | **1078.3** | **6.82** | **0.014** |
| GLMM 15 | 7 | 1078.5 | 7.06 | 0.012 |
| GLMM 20 | 11 | 1079.7 | 8.21 | 0.007 |
| GLMM 1 | 6 | 1080.5 | 9.05 | 0.005 |
| GLMM 0 | 4 | 1080.6 | 9.13 | 0.004 |
| GLMM 13 | 7 | 1081.7 | 10.23 | 0.003 |
| GLMM 10 | 9 | 1082.0 | 10.47 | 0.002 |
| GLMM 5 | 5 | 1082.0 | 10.52 | 0.002 |
| GLMM 14 | 7 | 1082.5 | 10.98 | 0.002 |
| GLMM 8 | 8 | 1082.5 | 11.04 | 0.002 |
| GLMM 2 | 6 | 1082.6 | 11.11 | 0.002 |
| GLMM 3 | 10 | 1085.2 | 13.75 | 0.000 |

**Table 3.**
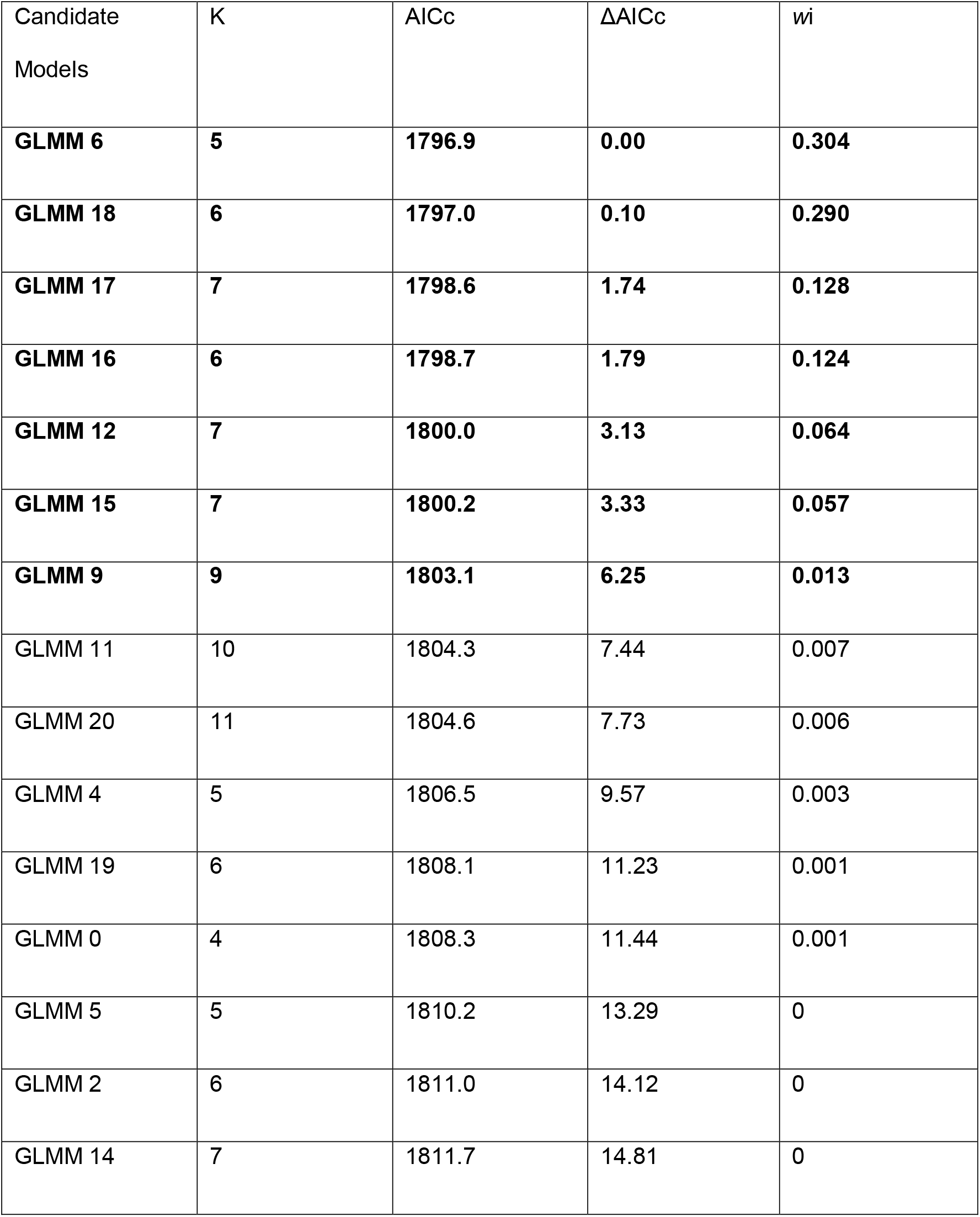
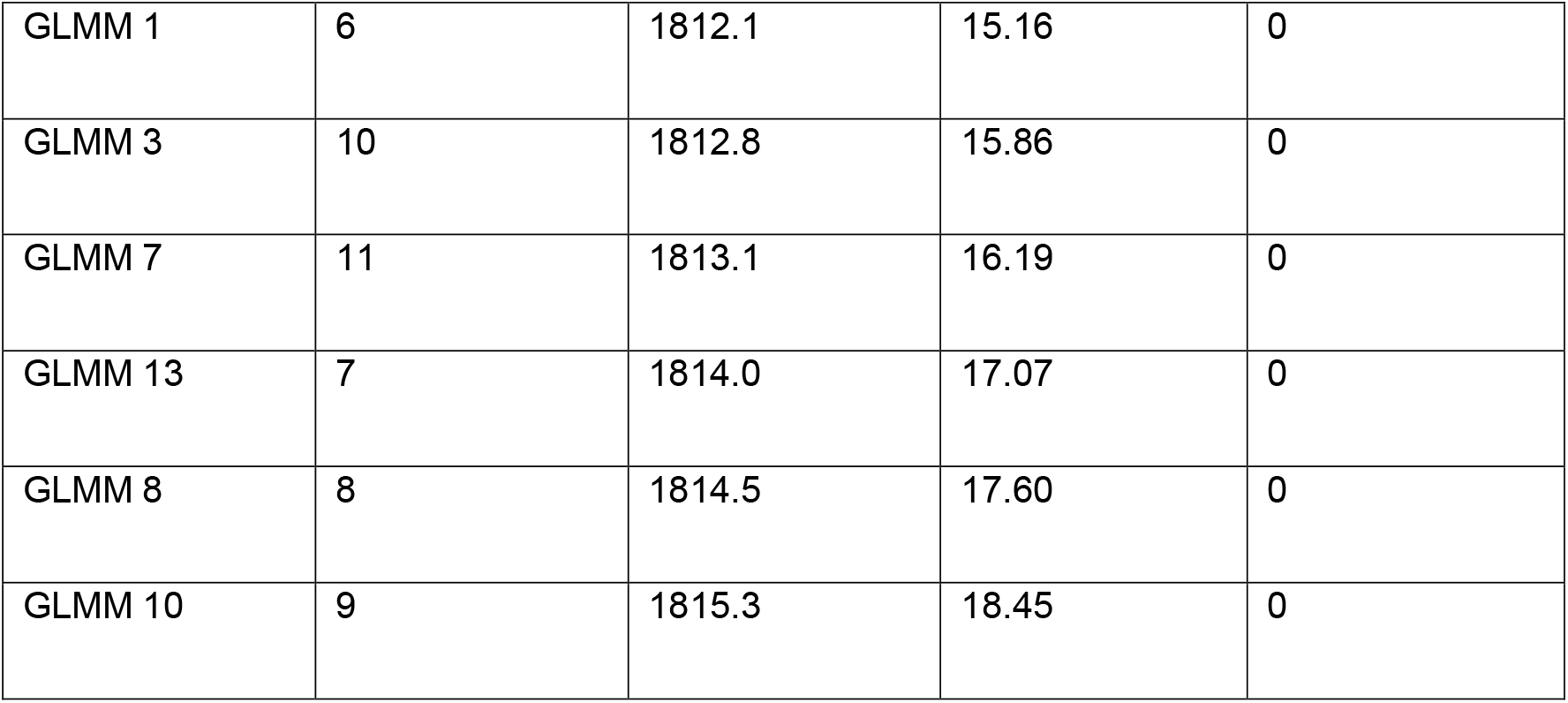
Summary of model-selection results for models explaining variation in SLEV exposure risk. K is the number of estimated parameters. Models are listed in decreasing order of importance. Models selected for model-averaged with ΔAICc > 7 are in bold.

| Candidate<br>Models | K | AICc | $\Delta AICc$ | $w_i$ |
| --- | --- | --- | --- | --- |
| **GLMM 6** | **5** | **1796.9** | **0.00** | **0.304** |
| **GLMM 18** | **6** | **1797.0** | **0.10** | **0.290** |
| **GLMM 17** | **7** | **1798.6** | **1.74** | **0.128** |
| **GLMM 16** | **6** | **1798.7** | **1.79** | **0.124** |
| **GLMM 12** | **7** | **1800.0** | **3.13** | **0.064** |
| **GLMM 15** | **7** | **1800.2** | **3.33** | **0.057** |
| **GLMM 9** | **9** | **1803.1** | **6.25** | **0.013** |
| GLMM 11 | 10 | 1804.3 | 7.44 | 0.007 |
| GLMM 20 | 11 | 1804.6 | 7.73 | 0.006 |
| GLMM 4 | 5 | 1806.5 | 9.57 | 0.003 |
| GLMM 19 | 6 | 1808.1 | 11.23 | 0.001 |
| GLMM 0 | 4 | 1808.3 | 11.44 | 0.001 |
| GLMM 5 | 5 | 1810.2 | 13.29 | 0 |
| GLMM 2 | 6 | 1811.0 | 14.12 | 0 |
| GLMM 14 | 7 | 1811.7 | 14.81 | 0 |
| GLMM 1 | 6 | 1812.1 | 15.16 | 0 |
| GLMM 3 | 10 | 1812.8 | 15.86 | 0 |
| GLMM 7 | 11 | 1813.1 | 16.19 | 0 |
| GLMM 13 | 7 | 1814.0 | 17.07 | 0 |
| GLMM 8 | 8 | 1814.5 | 17.60 | 0 |
| GLMM 10 | 9 | 1815.3 | 18.45 | 0 |

## Discussion

In this study, we analysed an extensive serological dataset on WNV and SLEV in relation to ecological and life-history traits of 132 bird’s species from different provinces in Argentina. We tested if selected ecological and life history traits could predict bird’s exposure risk to the afore mentioned viruses. Previous studies observed that nest characteristics and diet type influence the infection risk to mosquito-borne hemoparasites (González et al. 2014; Ganser et al. 2020), and that the number of offspring (i.e clutch size) and diet type positively correlate with other arboviral infections (Walsh 2019; Skinner et al. 2021). In addition, nest characteristics were previously associated with WNV infection risk in Senegal (Chevalier et al. 2009). Our study does not support the previously observed effects over WNV and SLEV exposure risk on the tested avian hosts species. Moreover, we observed that body mass and migratory status are important predictors for exposure risks to these arboviruses in our region. It should be noted that, differences in the selected traits between regions might be different due to changes in the maintenance networks of these mosquito-borne pathogens, particularly on those in which species of mosquitoes are involved in their transmission. We specifically observed that WNV exposure risk was higher in resident birds and increases as body mass increase. For SLEV, body mass was the only explanatory variable that predicted the exposure risk, being positively associated.

Previous studies have shown that body mass may play a significant role in vector-host contact rates, and hence, in host exposure risk to arbovirus (Takken & Verhulst 2013; Yan et al. 2017, 2018). The same pattern was observed for Ross River virus (a mosquito-borne arbovirus of the Togaviridae family) (Skinner et al. 2021), the vector-borne avian haematozoan parasites (Arriero & Møller 2008) and WNV in the southwest of Spain (Figuerola et al. 2008). Species with larger bodies, in which lead to higher body mass, might be more exposed to WNV and SLEV due the production of higher quantity of visual and olfactory cues, like carbon dioxide (Gillies & Wilkes 1972), and/or a diminished expression of anti-mosquitoes behaviours (Edman & Scott 1987; Mooring et al. 2004). Furthermore, they tend to live longer periods than smaller birds, thus increasing the likelihood to be exposed along their life span. Studies performed in Cx. pipiens mosquitoes showed that body mass affect mosquitoes’ feeding rates and host selection, even at an intraspecific level (Yan et al. 2017, 2018). This might indicate that both, intra and interspecific differences regarding ecological and life history traits are capable of modulating birds’ exposure risk by influencing mosquitoes feeding patterns and behaviour.

Migratory behaviour was an important predictor of WNV exposure risk, being resident species more exposed than migrants. Twenty six of the 132 sampled species catalogued as migrant exhibited 5 times lower exposure risk to WNV. Within these species, both Neotropical and Nearctic long-distance migrants were included (Hayes 1995). The former breeding in Argentina and migrating northward during non-breeding season, like the Vermilion flycatcher (Pyrocephalus rubinus), the Double-collared seedeater (Sporophila caerulescens), the White-banded Mockingbird (Mimus triurus) and species of the Elaenia genera. The latter, Nearctic long-distance migrants, breeds in North America and migrates southward during non-breeding season, like the White-rumped sandpiper (Calidris fuscicollis) and the Stilt sandpiper (Calidris himantopus). Despite these species show differences in their migratory behaviour, in addition to differences in other ecological and life history traits, none of them showed to be exposed to WNV nor SLEV (seroprevalence ranging from 0.007 to 0.04). These could indicate that neither the migratory routes nor the migratory behaviour of the evaluated species affects their exposure risk differentially. The observed values in exposure risk could be due to differences in viral circulation determined by local variations in the transmission networks between breeding and non-breeding areas (Kilpatrick et al. 2006b; Diaz et al. 2013b), as observed in Palearctic summer migrants in Spain and resident birds in Senegal (Figuerola et al. 2008; Chevalier et al. 2009). Lastly, it is important to highlight that exposure risk does not reflect the potential role of a specie in the transmission network, as it has been largely proposed that migratory birds are partially responsible in viral spreading (Rappole et al. 2000; Hoover & Barker 2016; Swetnam et al. 2018).

Life history traits are not only related with birds’ exposure risk to vector-borne pathogens by modulating host-vector encounters, but they are also closely related to birds’ immune function throughout the pace of life syndrome and trade-offs (Norris & Evans 2000; Zuk & Stoehr 2002; Tieleman 2018). Within this context, species with “slow-living” traits (longer life cycles, larger body size, slower reproductive rates, and higher adult survival) show differences in immune capacity and responses in contrast with “fast-living” species (characterized by opposite traits in the pace of life spectrum) (Martin et al. 2006; Hasselquist 2007; Johnson et al. 2012). Particularly, slow-living species will tend to favour specific immune defences over constitutive and proinflammatory responses (Lee 2006; Martin et al. 2006, 2008). On the other hand, “fast-living” species will tend to emphasise more in the unspecific components of their immune responses (Lee 2006; Lee et al. 2008a). Moreover, life history traits like long-distance migrations are linked to trade-offs as well, limiting resources that could be used in immune function and, hence (Schmid-Hempel & Ebert 2003), modifying birds resource allocation (Buehler et al. 2010; Hasselquist & Nilsson 2012).

Traits like body mass and clutch size (that are included in the pace of life syndrome) has been previously used to explain differences in host competence index for WNV and Eastern Equine Encephalitis Virus, being negatively correlated with body mass and positively with clutch size (Huang et al. 2013). Now, knowing which life history traits were better able to predict WNV and SLEV exposure risks, we might ask if these selected traits can provide information about the possible outcome of viral infection. Both body mass and migration status have been shown to be related with the immune response. Firstly, body mass was used to explain differences in host competence index for WNV and Equine Encephalitis Virus (Huang et al. 2013). This pattern might be due to the relationship established between body mass, the pace of life continuum and immune function (Lee 2006). Specifically, larger slow-living animals develop stronger induced cellular immune responses and relay more heavily on developing induced humoral immune responses, which are less expensive from an energetic perspective (Lee 2006; Palacios & Martin 2006; Lee et al. 2008b; Rauw 2012). These traits might give advantages to larger birds against their greater exposure risk when responding to arboviral infection while facing multiple environmental demands.

On the other hand, and regarding the differences between migrant and resident birds, migrants might compromise their ecological capital and resource allocation towards migration as a priority (Zuk & Stoehr 2002; Lee & Klasing 2004; Buehler et al. 2010). Consequently, they are not expected to have the full capacity to mount the required immune response against viral infection (Sitati et al. 2003; Rauw 2012). Nevertheless, a study performed by Owen et al. showed that there were not significant effects of displaying migratory activity over the infection outcome, but affected their migratory activity, changing the status of half the cohort to non-migratory (Owen et al. 2006). This might indicate that if the infection is prior to the migratory event, the resource allocation will favour solving the infection at the expense of migration. Still, it is uncertain what the outcome will be if the infections occur temporarily after the migratory event. In this context, it is possible for species with higher exposure risks (resident and larger) to have the resources to resist viral infection, thus being capable to afford the costs of the complex immune response required to resist viral infection (Zuk & Stoehr 2002; Samuel & Diamond 2006; Rauw 2012). Hence, these species might exhibit a lower contribution to the viral flow in the arboviral maintenance network, highlighting the fact that the trade-offs in viral amplification driven by immune system are also needed to be considered in transmission dynamics (Althouse & Hanley 2015).

Overall, our results suggests that body mass is the main driver or birds’ exposure risk to WNV and SLEV. Moreover, larger resident birds might be prone to WNV exposure compared to smaller migrant birds. These results also provide information on our regional transmission system of these arboviruses, being useful to focus resources when describing local viral prevalence and characterizing viral transmission networks. Lastly and based on an immune-ecological approach, the most exposed bird’s species might not be efficient host for the evaluated viruses, thus compensating the potential increased in transmission. This theoretical approach should be carefully considered although it needs to be further explored experimentally, considering the trade-offs that might occur within host viral replication.

## Acknowledgments

We are grateful to M.D. Beranek, A. Visintin, M. Laurito, M. Stein, G. Oria, J. Rosas, MJ Dantur Juri, R. Lobos, E. Seiler, S. Flores and Batallán, PG for their technical assistance during sample collection and serological determinations. We also thanks to A. I. E. Quaglia for his assistance in statistical analysis and data visualization. This project was supported by grants PICT 2018-01172, SECYT Consolidar UNC and PUE 2016 IIBYT-CONICET. Octavio Giayetto is a PhD candidate and recipient of doctoral scholarship from CONICET.

## Data, scripts, code, and supplementary information availability

Data and R-codes are available in figshare repository (Giayetto et al. 2023; DOI: https://doi.org/10.6084/m9.figshare.21978875.v1). The method section and description provide all necessary information about the dataset.

## Statement of authorship

GO, NFN and DA designed the research, GO collected data, GO and MAP performed the statistical analysis, all authors analysed the output data. GO wrote the first draft of the manuscript and all authors contributed with the manuscript revision.

